# Longitudinal transcriptomic analysis reveals adaptive transcript usage remodeling associated with PARP inhibitor resistance

**DOI:** 10.64898/2026.09.22.753421

**Authors:** Jiye Yoo, Yoo-Na Kim, Yewon Jung, Eunhae Cho, Jung Kyoon Choi, Hyeon Gu Kang, Jung-Yun Lee

## Abstract

Poly (ADP-ribose) polymerase inhibitors (PARPi) have revolutionized the treatment of ovarian and breast cancers with homologous recombination deficiency (HRD). Despite substantial initial efficacy, the frequent development of acquired resistance poses a critical clinical challenge, necessitating a deeper understanding of the underlying molecular mechanisms. While genomic profiling provides information on baseline HRD status, it is insufficient to capture the dynamic and functional adaptations accompanying resistance during treatment.

In this study, we investigated the role of alternative splicing in PARPi resistance using longitudinal pre- and post-treatment RNA-sequencing data from an ovarian cancer patient cohort (n=40). We demonstrated that profiling alternative splicing dynamics and differentially used transcripts (DUTs) provides transcriptomic insights distinct from conventional differential gene expression analysis. Notably, we observed significant differences between acquired resistance (AR) and innate resistance (IR) groups. Specifically, longitudinal analysis revealed that AR tumors exhibit a distinct transcript usage pattern, characterized by an increase in canonical protein-coding transcripts involved in cell-cycle processes and a decrease in non-canonical non-coding transcripts within the homologous recombination pathway following treatment. Extending this comparison across AR and IR tumors at both pre- and post-treatment stages, these transcriptomic features were most pronounced in the AR post-treatment group. Further analysis of subsequent treatment outcomes showed that, among AR patients, greater resistance-associated transcriptomic remodeling was associated with poorer response and shorter progression-free survival. The resistance-associated transcriptomic signatures identified through longitudinal profiling were further recapitulated in an independent pre-treatment cohort (n=71; 43 AR and 28 IR), with more pronounced differences among BRCA wild-type patients. Taken together, our findings identify differential transcript usage as dynamic transcriptomic features associated with acquired PARPi resistance and establish a framework for utilizing longitudinal transcriptome dynamics to investigate the molecular basis of clinical drug resistance.

## Introduction

Poly (ADP-ribose) polymerase (PARP) plays a critical role in sensing single strand breaks (SSBs) and coordinating their repair by recruiting downstream repair machinery^1,2^. Thus, PARP maintains genomic stability and ensures cell survival under genotoxic stress. Capitalizing on the concept of synthetic lethality, PARP inhibitors (PARPi) have demonstrated profound cytotoxicity in cancer cells with impaired *BRCA1* and *BRCA2* function^3,4^. Consequently, various genomic profiling methods–such as quantifying structural variations (SVs), single-nucleotide variants (SNVs), and mutational signatures^5–9^–have been implemented to evaluate baseline HRD status and predict clinical responses to PARPi. Additionally, various methods are being studied that focus on several core genes of the DNA repair pathway as biomarkers, such as *BRCA1*, *BRCA2*, *RAD51*, and *ATM*^10^. However, static genomic biomarkers possess inherent limitations; they cannot capture the real-time functional status of DNA repair pathways or track dynamic adaptations during treatment. This limitation makes it difficult to detect or predict acquired resistance, a major clinical challenge that contributes to disease recurrence in advanced-stage ovarian cancer patients receiving PARPi and platinum-based therapies, with recurrence occurring in up to 80% of patients^11–15^. While several resistance mechanisms have been documented–including secondary reversion mutations that restore BRCA1/2 function, upregulation of drug efflux pumps, and epigenetic remodeling16–genomic alterations alone do not fully explain the emergence of clinical resistance, suggesting that non-genomic mechanisms may contribute to PARPi resistance. Alternative splicing is a major post-transcriptional regulatory mechanism that expands transcriptomic and proteomic diversity by generating multiple transcript isoforms from a single gene through differential exon and splice-site selection^17,18^. These isoforms can differ in coding potential, subcellular location, RNA stability, and functional domain composition, thereby enabling individual genes to produce functionally distinct transcripts in response to specific biological contexts and environmental stresses^19–24^. Consequently, dysregulation of RNA splicing can profoundly alter cellular function and has been implicated in a wide range of human diseases, including cancer and genetic disorders^23,25,26^. Thus, changes in transcript isoform composition can represent an important layer of cellular adaptation, particularly under selective pressures such as anticancer therapy. However, despite the potential role of transcriptomic remodeling in therapeutic adaptation, previous studies have largely characterized PARPi-associated genomic features and gene-level transcriptional changes in pre-treatment patient samples or cell-line models, while how transcriptomic programs change longitudinally during acquired resistance in patient tumors remains poorly understood^27,28^. Building upon our previous findings that baseline transcriptome configurations can serve as predictive markers for PARP inhibitor response^29^, we extended our investigation to longitudinal samples to examine how transcript usage changes during treatment and whether these changes are associated with acquired resistance.

In this study, we investigated the dynamic changes in alternative splicing and transcript usage by analyzing longitudinal, paired ovarian cancer samples collected before PARPi treatment and at recurrence in patients with acquired-resistance (AR) or innate-resistance (IR) groups. We demonstrated that analyzing differentially used transcripts (DUTs) uncovers transcript-level adaptations that are not readily detected by standard differential gene expression analysis. Notably, we discovered significant longitudinal differences between the AR and IR cohorts. Within the AR group, post-treatment samples exhibited a distinct transcript switching pattern characterized by an increase in functional transcripts coding for cell-cycle process, coupled with a concomitant decrease in non-functional transcripts within the homologous recombination (HR) pathway. By evaluating these dynamic transcript configurations, we found that the post-treatment AR group exhibited the strongest transcriptomic signature of DNA repair activation among all examined AR and IR subsets. Furthermore, our analysis suggested that this resistance-associated transcriptomic signature could stratify PARPi responsiveness—significantly influenced by the patients’ BRCA mutation status—and showed that this signature was reproducible in an independent, external pre-treatment clinical cohort. Taken together, our findings provide a comprehensive look into how alternative splicing serves as a dynamic non-genomic resistance mechanism under therapeutic pressure, highlighting the clinical value of longitudinal isoform tracking to predict and monitor PARPi resistance in ovarian cancer.

## Results

### Study Design and Clinical Characteristics of the Longitudinal PARPi Cohort

To understand how dynamic molecular changes contribute to PARP inhibitor (PARPi) resistance, we performed longitudinal RNA-sequencing on ovarian cancer patients before (pre-treatment; baseline) and after (post-treatment; relapse) PARPi treatment (n=40). Based on clinical responses and recurrence timelines, patients were classified into two distinct groups: acquired resistance (AR, n=23) and innate resistance (IR, n=17) (**Fig. 1a** and **Methods**). The AR group comprised patients who initially derived clinical benefit from PARPi treatment but subsequently relapsed over time. In contrast, the IR group consisted of patients who exhibited intrinsic resistance without an initial response to PARPi. This well-defined longitudinal cohort provides a clinical framework to track and compare the molecular adaptations driving acquired versus innate resistance.

**Figure 1.**
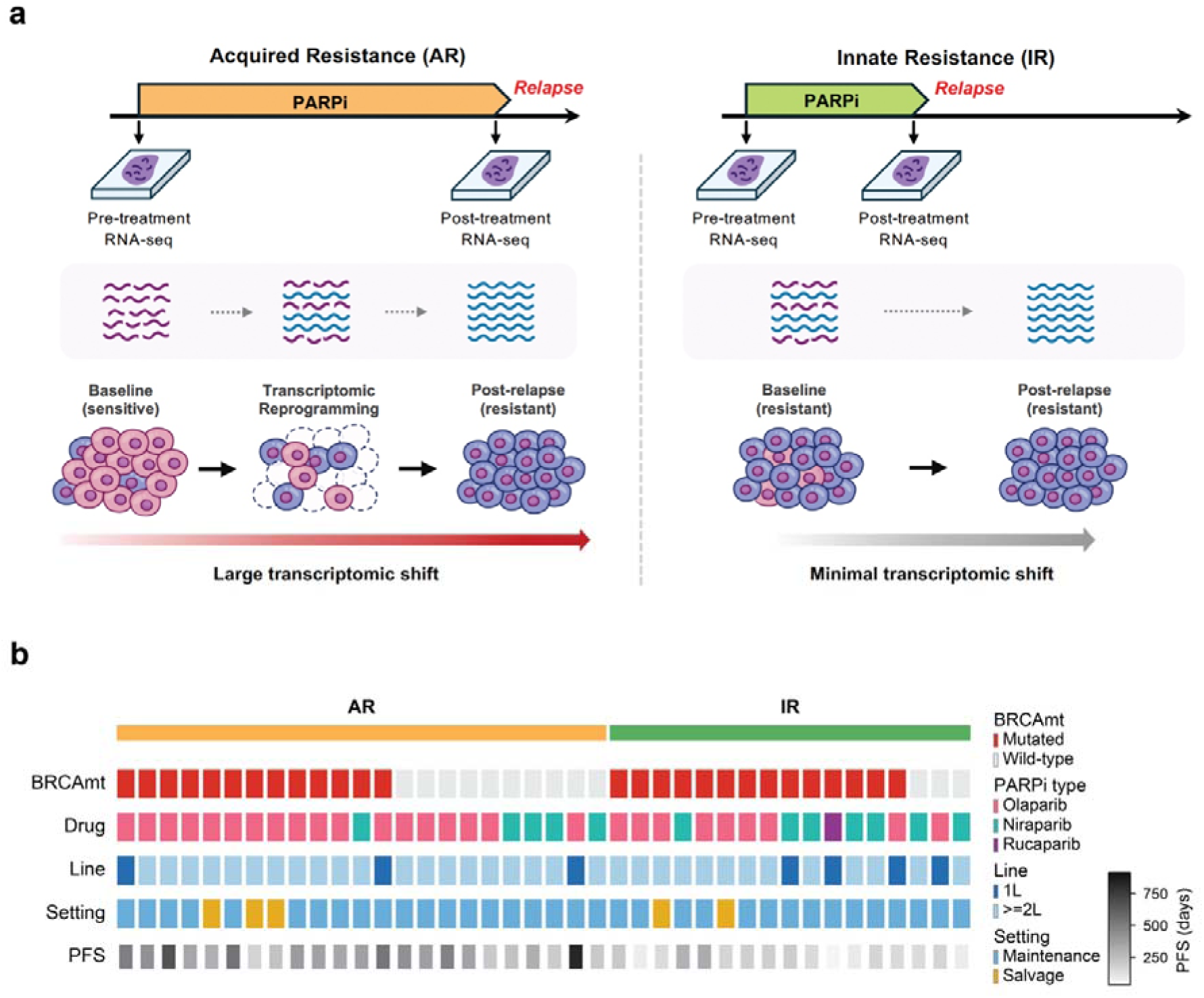
Overview of the longitudinal PARPi-treated ovarian cancer cohort. (a) Schematic overview of the longitudinal study design. Paired tumor samples were collected at pre-treatment and post-treatment relapse from patients with acquired resistance (AR) or innate resistance (IR) to PARP inhibitors (PARPi) and subjected to RNA sequencing. (b) Clinical characteristics of patients in the longitudinal cohort. Each column represents an individual patient, grouped by AR and IR status. Clinical annotations include BRCA mutation status (BRCAmt), treatment line (1L or ≥2L), administered PARPi types (olaparib, niraparib, or rucaparib), therapeutic setting (maintenance or salvage), and progression-free survival (PFS, days).

To minimize potential clinical confounding, we first characterized the clinical features of our longitudinal cohort (**Fig. 1b**). These included BRCA mutation status (BRCA mutant vs. wild-type), treatment line (1L vs ≥2L), administered PARPi types (olaparib, niraparib, and rucaparib), therapeutic setting (maintenance vs. salvage), and progression-free survival (PFS, days). Importantly, no statistically significant differences were observed between the AR and IR groups across these clinical features, indicating that the groups were generally comparable across the evaluated clinical variables (**Supplementary Table 1**). These findings suggest that the two groups were clinically comparable at baseline, providing an appropriate framework for comparing longitudinal transcriptomic changes while reducing the likelihood that the observed differences are explained by major clinical features.

### Widespread Transcriptomic Reprogramming Characterizes Acquired PARPi Resistance

We next investigated the specific molecular changes underlying the divergent resistance phenotypes. Differential expression analysis identified substantially more treatment-associated changes in AR than in IR tumors, with 1,918 DEGs in AR compared with 425 in IR (**Fig. 2a**). Functional enrichment analysis of AR-associated DEGs revealed significant enrichment of DNA repair-related processes, whereas no significantly enriched pathways were identified among IR-associated DEGs (**Supplementary Fig. 1**). Despite this pathway-level enrichment, genes directly involved in homologous recombination (HR)^30,31^, including *BRCA1/2*, *RAD51*, and *ATM*, showed relatively limited changes in overall gene expression. This lack of direct gene-level changes prompted us to analyze alternative splicing (AS) which expands transcriptomic diversity by producing multiple isoforms from a single gene. Notably, alternative splicing analysis demonstrated a substantial elevation in splicing events— including retained introns (RI), skipped exons (SE), and alternative 3’/5’ splice sites (A3SS/A5SS)—specifically in AR tumors (**Fig. 2b** and **Methods**). Consistent with our previous DEG analysis, this prominent bias reinforces the notion that AR tumors selectively undergo significant transcriptomic reshaping between pre- and post-treatment states. To explore potential regulators of these widespread splicing changes, we compared splicing factor (SF) expression between pre- and post-treatment tumors (**Methods**). Splicing factor genes showed significant expression changes exclusively in AR tumors (**Fig. 2c**) with key individual SFs such as *DDX1, PRPF6, HNRNPF,* and *RBM47* significantly upregulated post-treatment (**Fig. 2d**). This coordinated pattern suggests increased splicing activity specifically in AR tumors. We next examined the magnitude and direction of these splicing changes using the distribution of ΔPSI (percent spliced in). The ΔPSI distribution for RI events revealed a pronounced directional difference between AR and IR tumors (**Fig. 2e**). Specifically, negative ΔPSI RI events were significantly more frequent in AR than in IR tumors (OR = 2.22, two-sided Fisher’s exact p = 8.0 × 10^−9^), indicating a stronger shift toward reduced intron retention following treatment in AR tumors. This pattern is consistent with reduced production of transcripts susceptible to nonsense-mediated decay (NMD), suggesting that alternative splicing may contribute to preserving functional transcripts. Together, these findings indicate that transcriptomic remodeling through AS is substantially more pronounced in AR than IR tumors following PARPi treatment.

**Figure 2.**
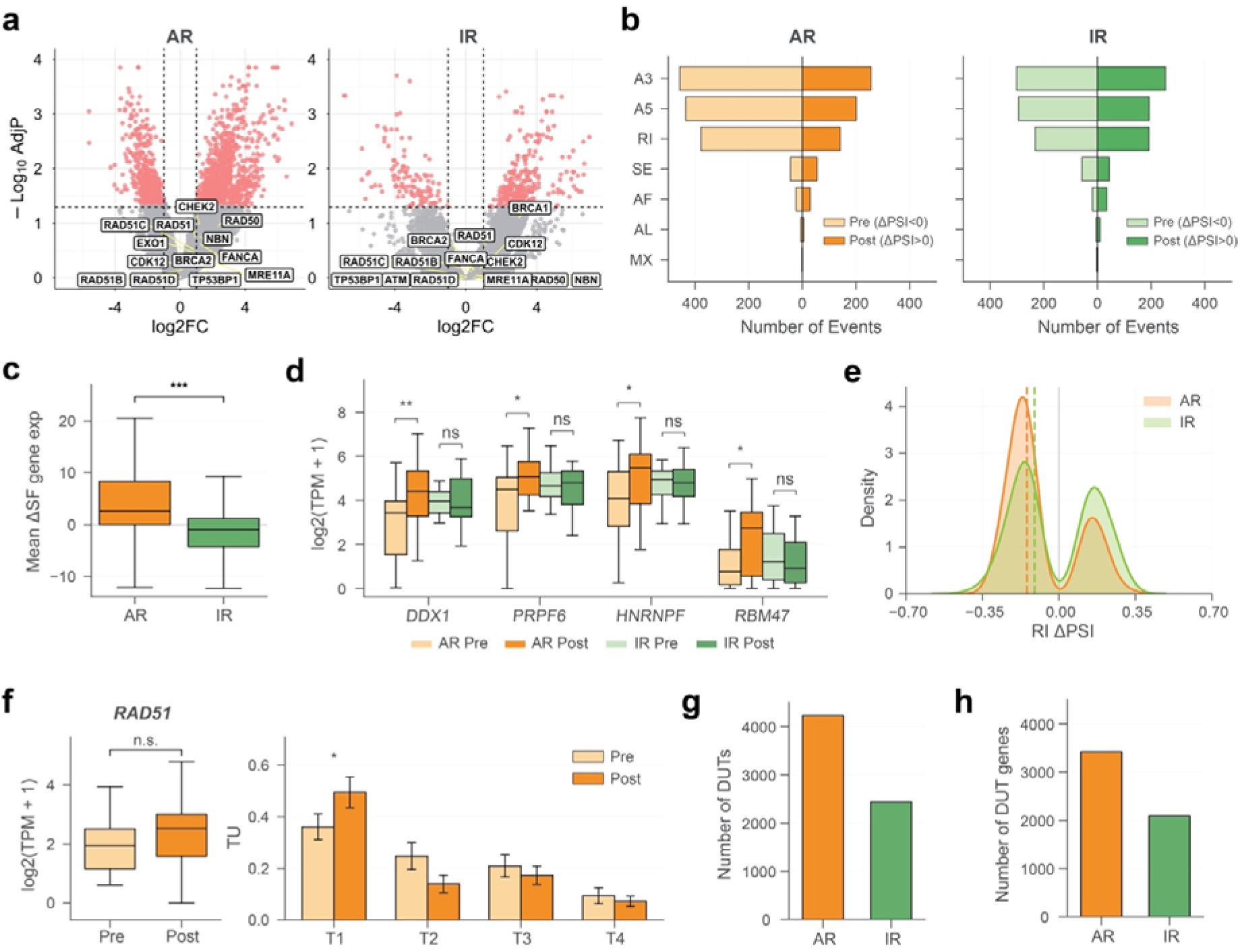
Multi-level transcriptomic changes in acquired and innate PARPi resistance. (a) Differential gene expression (DEG) volcano plots comparing pre- and post-treatment tumors within the AR and IR groups. Genes previously implicated in PARPi response are highlighted. (b) Number of significant alternative splicing events identified in AR and IR samples, stratified by the direction of ΔPSI (post - pre). Across event types, AR tumors showed 713 A3, 636 A5, 519 RI, 100 SE, 53 AF, 11 AL, and 3 MX events, whereas IR tumors showed 555 A3, 487 A5, 424 RI, 102 SE, 54 AF, 15 AL, and 2 MX events. “Pre” and “Post” indicate negative and positive ΔPSI (post - pre), respectively. (c) Mean change in splicing factor gene expression following treatment in AR and IR tumors. P values were calculated using two-sided Mann-Whitney U tests. (d) Expression changes of representative splicing factor genes in AR and IR tumors. P values were calculated using two-sided Wilcoxon signed-rank tests. (e) Density distributions of ΔPSI values for retained intron (RI) events in AR and IR tumors. (f) Representative example of transcript usage changes in RAD51, showing total gene expression and transcript-level TU between pre- and post-treatment AR tumors. T1–T4 denote individual RAD51 transcript isoforms. P values were calculated using two-sided Wilcoxon signed-rank tests. (g) Number of DUTs in AR (n=4,231) and IR (n=2,434) samples. (h) Number of genes harboring DUTs in AR (n=3,422) and IR (n=2,087) samples.

Because individual splicing events provide limited information on the resulting transcript isoforms and their functional properties, we next evaluated transcript usage (TU) to capture isoform-level dynamics (**Methods**). For instance, analysis of *RAD51* in AR tumors showed no significant difference in total gene expression between pre- and post-treatment states (**Fig. 2f**, **left panel**), whereas its transcript usage exhibited a significant shift following treatment (**Fig. 2f**, **right panel**), enabling transcript-level characterization of the functional consequences of splicing changes. Consistent with this observation, several additional PARPi-related genes showed differential transcript usage despite the absence of significant gene-level expression changes (**Supplementary Fig. 2**), further highlighting the distinct information captured by isoform-level analysis. We reasoned that integrating both quantitative and qualitative analyses would provide an additional insight to understand the resistance mechanisms triggered by transcriptomic changes. Consequently, we performed statistical testing on TU alterations between pre- and post-treatment samples for each group (**Methods**), uncovering a greater number of differentially used transcripts (DUTs) and genes harboring DUTs in the AR tumors than in the IR tumors (**Fig. 2g** and **h**). Together, these findings demonstrate that acquired resistance is characterized by active transcriptomic changes rather than a static resistance state.

### Functional Classification of AR-Associated DUTs Reveals Transcriptomic Remodeling of Cell-Cycle and Homologous Recombination Pathways

To elucidate the functional impact of the identified DUTs in AR tumors (**Fig. 3a, left panel**), we categorized AR DUTs according to their resistance-associated direction of change. Transcripts with decreased TU post-treatment were designated as PSTs (PARPi-sensitivity transcripts, n=723), while those with increased TU were defined as PRTs (PARPi-resistance transcripts, n=1,344) (**Fig. 3a, middle panel**). We next considered their functional features and further classified these transcripts as Class 1 (canonical protein-coding), Class 2 (non-canonical protein-coding), and Class 3 (non-canonical non-coding) (**Fig. 3a, right panel** and **Methods**). Due to the functional ambiguity of Class 2 transcripts, only Class 1 and Class 3 transcripts were included in subsequent analyses.

**Figure 3.**
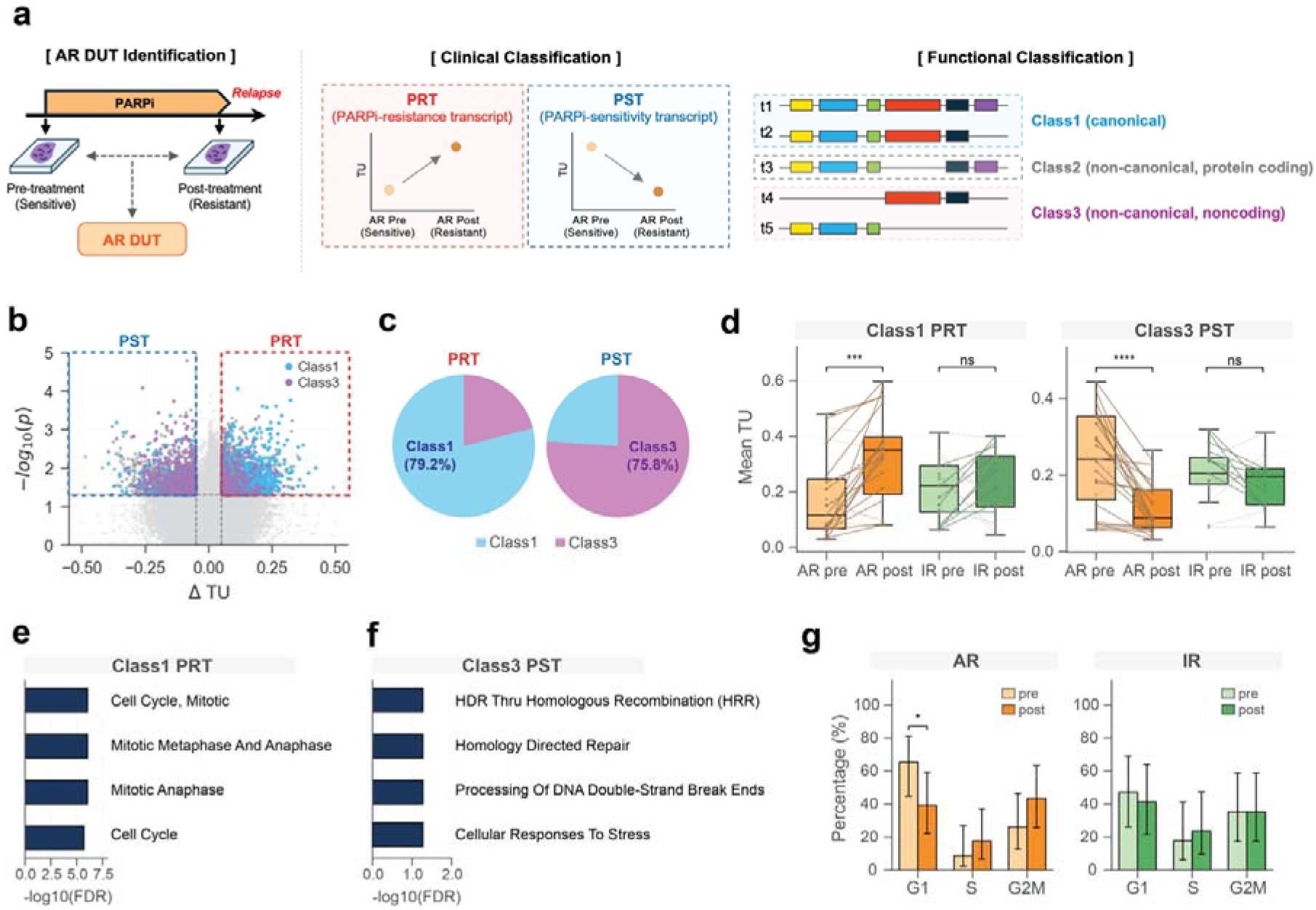
Functional classification and pathway-specific characteristics of differentially used transcripts associated with acquired PARPi resistance. (a) Schematic diagram of clinical classification and functional classification of AR DUTs. (b) Volcano plot of transcript usage changes in AR tumors, highlighting Class 1 and Class 3 transcripts among PRTs and PSTs. (c) Proportions of Class 1 and Class 3 transcripts among PRTs and PSTs. (d) Mean transcript usage of Class 1 PRTs and Class 3 PSTs across the four clinical groups. Solid lines indicate patients showing resistance-consistent changes (increased Class 1 PRT usage or decreased Class 3 PST usage post-treatment), whereas dashed lines indicate all other patients. P values were calculated using two-sided Wilcoxon signed-rank tests. (e) Top enriched pathways among Class 1 PRT genes (n=1,055). (f) Top enriched pathways among Class 3 PST genes (n=540). (g) Cell-cycle phase distributions (G1, S, and G2/M) in pre- and post-treatment AR and IR tumors. P values were calculated using two-sided Wilcoxon signed-rank tests for paired pre- and post-treatment comparisons.

Intriguingly, 1,064 PRTs (79.2%) were classified as Class 1 canonical protein-coding transcripts, whereas 548 PSTs (75.8%) were classified as Class 3 non-coding transcripts (**Fig. 3b** and **c**). To ensure that these transcriptomic signatures were not driven by a subset of outlier samples, we examined the paired pre-versus post-treatment changes at the individual sample level within both the AR and IR groups. Strikingly, these distinct transcriptomic signatures (Class 1 PRTs and Class 3 PSTs) showed consistent shifts in the expected directions following treatment in the AR group, whereas no significant changes were observed in the IR group (**Fig. 3d**). These findings further support the association of these transcriptomic changes with the acquired resistance phenotype. At the individual-patient level, however, the specific transcripts contributing to these shifts varied substantially across AR tumors, with each patient exhibiting a distinct combination of increased usage of Class 1 PRTs and decreased Class 3 PSTs (**Supplementary Fig. 3a** and **b**). Despite the heterogeneity of individual transcript changes, the significant difference in coding potential between PRTs and PSTs suggested distinct biological roles for the two signatures. We therefore performed functional enrichment analysis to identify the pathways associated with each signature. Class 1 PRTs were strongly enriched in pathways regulating the mitotic cell cycle and anaphase progression^12,32^ (**Fig. 3e**), while Class 3 PSTs were enriched in homology-directed repair (HDR) and DNA double-strand break processing (**Fig. 3f**). These results suggest that PARPi resistance is associated with adaptive isoform switching that remodels transcript composition in pathways involved in cell-cycle regulation and homologous recombination repair, reflecting potential transcriptomic adaptation to PARPi treatment. To assess whether these transcriptomic signatures were accompanied by corresponding cellular phenotypes, we examined the cell cycle distribution across the groups (**Methods**). In line with the Class 1 PRT enrichment result, AR tumors displayed a significant reduction in the G1 phase population and an increased G2/M trend post-treatment, a phenotypic shift not observed in IR tumors (**Fig. 3g**). Together, these findings link coordinated increases in Class 1 PRT usage and decreases in Class 3 PST usage to transcriptomic programs associated with cell-cycle progression and homologous recombination repair. Hereafter, the PRT and PST signatures are defined as the mean TU of Class 1 PRTs and Class 3 PSTs, respectively, within each sample.

### Patient-Level Transcriptomic Signatures Characterize PARPi Resistance States and Subsequent Treatment Outcomes

Beyond the pairwise comparison of pre- and post-treatment changes within each group, we asked whether the transcriptomic signature changes observed during acquired resistance could be identified in intrinsically resistant IR-pre tumors. Integrating all four clinical subsets revealed a progressive, stepwise trend in these transcriptomic signatures across AR-pre, IR-pre, IR-post, and AR-post tumors, suggesting a continuous trajectory toward PARPi resistance (**Fig. 4a** and **d**). We reasoned that this stepwise trend not only elucidates the underlying mechanism but could also reflect subtle baseline transcriptomic states that predispose tumors to differential therapeutic response. Principal component analysis (PCA) and cluster proportions based on these transcriptomic signatures further supported the existence of the stepwise trend. Rather than forming discrete transcriptomic states, the four clinical subsets were progressively ordered along the resistance axis. For the PRT signature, the clusters were ordered from AR-pre-enriched C1, through IR-enriched C0, to AR-post-enriched C2, whereas the PST signature showed an increased proportion of AR-post tumors from C0 to C1 (**Fig. 4b** and **e**). Projection of individual samples onto the IR-derived transcriptomic axis further confirmed that 74% of AR tumors shifted toward an IR-like state following treatment, highlighting a shared trajectory underlying both acquired and innate resistance (**Fig. 4c, f,** and **Methods**). We next examined whether this resistance-associated pattern was preserved within the cell-cycle and HRR pathways identified in our preceding functional enrichment analysis. Pathway-specific cell-cycle PRT and HRR PST signatures showed trends consistent with the global PRT/PST patterns (**Supplementary Fig. 4**).

**Figure 4.**
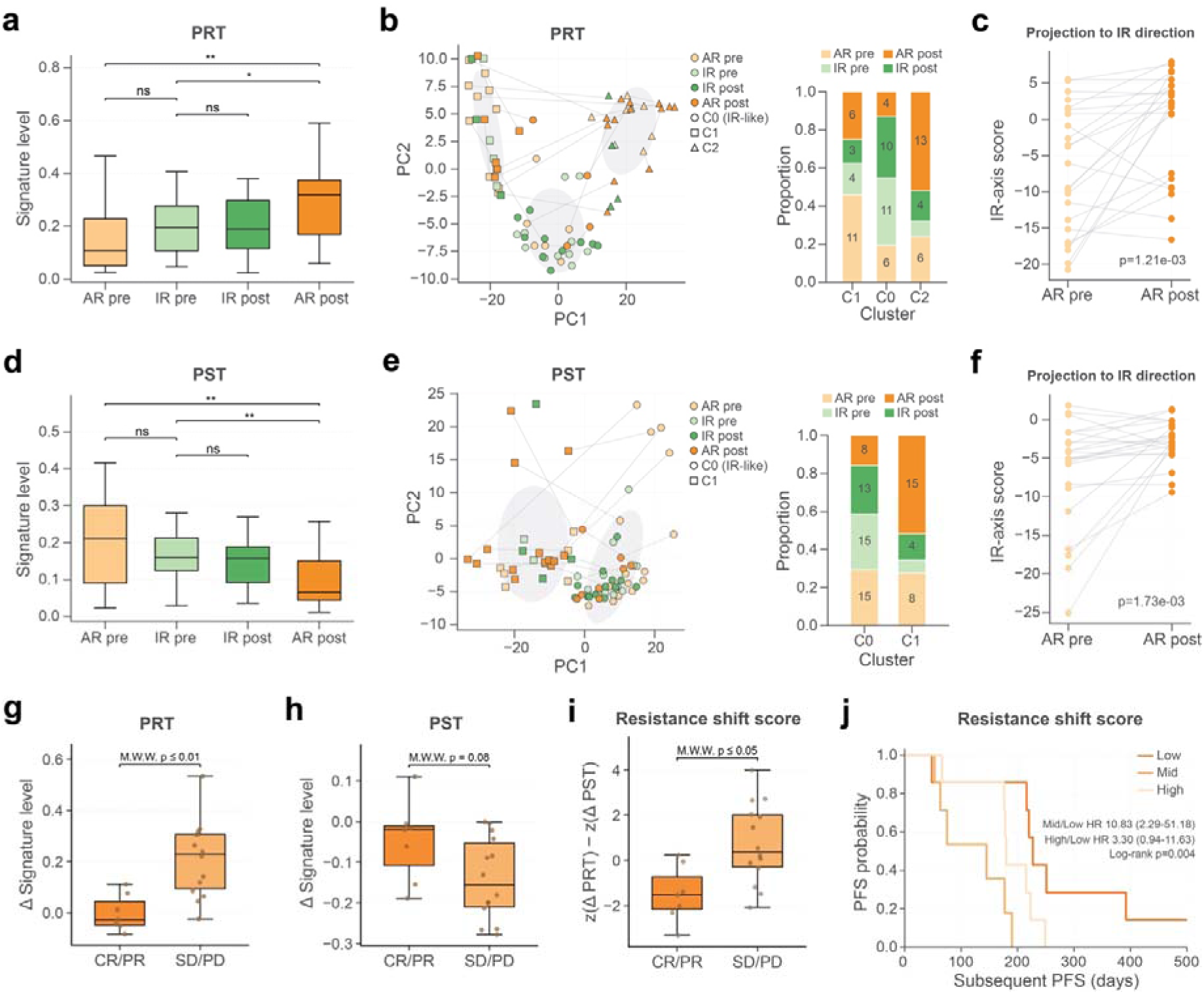
Resistance-associated transcriptomic signatures and their association with subsequent treatment outcomes. (a) PRT signature levels across the four clinical groups (AR pre, IR pre, IR post, and AR post). P values were calculated using two-sided Mann-Whitney U tests. (b) Principal component analysis (PCA) of samples based on PRT transcript usage. Samples are colored by clinical group and annotated according to cluster assignment. The stacked bar plot shows the composition of clinical groups within each cluster. (c) Projection of paired AR pre- and post-treatment samples onto the IR-derived PRT axis. P values were calculated using two-sided Wilcoxon signed-rank tests. (d) PST signature levels across the four clinical groups (AR pre, IR pre, IR post, and AR post). P values were calculated using two-sided Mann–Whitney U tests. (e) Principal component analysis (PCA) of samples based on PST transcript usage. Samples are colored by clinical group and annotated according to cluster assignment. The stacked bar plot shows the composition of clinical groups within each cluster. (f) Projection of paired AR pre- and post-treatment samples onto the IR-derived PST axis. P values were calculated using two-sided Wilcoxon signed-rank tests. (g) Changes in PRT signature levels during PARPi treatment according to subsequent treatment response among AR patients. Patients were grouped as CR/PR or SD/PD according to their response to subsequent therapy. P value was calculated using two-sided Mann-Whitney U tests. (h) Changes in PST signature levels during PARPi treatment according to subsequent treatment response. P value was calculated using two-sided Mann-Whitney U tests. (i) Resistance shift score according to subsequent treatment response. The score integrates opposing changes in the PRT and PST signatures and was calculated as z(ΔPRT) − z(ΔPST), where ΔPRT and ΔPST represent treatment-associated changes in the corresponding signature levels. P value was calculated using two-sided Mann-Whitney U tests. (j) Kaplan-Meier analysis of progression-free survival (PFS) following subsequent therapy according to tertiles of the resistance shift score. Patients were classified into low-, intermediate-, and high-shift groups.

To further assess the clinical relevance of this longitudinal remodeling within AR tumors, we analyzed subsequent treatment outcomes in 21 of the 23 AR patients for whom post-PARPi treatment information was available (**Supplementary Table 2**). AR patients who had stable disease (SD) or progressive disease (PD) to subsequent therapy exhibited significantly greater increases in PRT signature levels during PARPi treatment than those who achieved partial response (PR) or complete response (CR) (**Fig. 4g**). A concordant trend toward greater decreases in PST signature levels was also observed in the SD/PD group (**Fig. 4h**). Consistently, integration of the opposing changes in the PRT and PST signatures into a resistance shift score revealed significantly higher scores in the SD/PD group than in the PR/CR group (**Fig. 4i**). Furthermore, tertile-based stratification revealed that only patients with minimal resistance-associated transcriptomic remodeling showed prolonged subsequent PFS, whereas intermediate- and high-shift groups showed similarly shorter subsequent PFS (**Fig. 4j**). Notably, pathway-specific analysis revealed significant differences in both cell-cycle PRT and HRR PST signature shifts between the response groups, with greater increases in the cell-cycle PRT signature and decreases in the HRR PST signature in patients with SD/PD (**Supplementary Fig. 5**). The corresponding pathway-specific resistance shift scores also reproduced the subsequent treatment response and PFS patterns observed with the global resistance shift score. Together, these findings suggest that the magnitude of transcriptomic remodeling acquired during PARPi resistance is associated not only with PARPi resistance itself but also with poorer response and shorter PFS following subsequent therapy, potentially reflecting a broader treatment-refractory phenotype.

We next investigated whether these pathway-specific trends were also reflected at the individual gene level. Consistent with these observations, representative cell-cycle PRTs from *RAD51AP1* and *TOPBP1* and HRR PSTs from *RBBP8* and *RECQL* showed pronounced transcript usage changes in AR post-treatment tumors despite subtle changes in overall gene expression (**Supplementary Fig. 6**). These results further indicate that coordinated changes in the usage of functionally relevant transcripts, rather than broad changes in gene expression, accompany the transcriptomic transition toward PARPi resistance.

Together, these findings suggest that PARPi resistance is associated with ordered changes in transcriptomic states, with the magnitude of this remodeling also associated with response to subsequent therapy. The consistent progression observed across all four clinical subsets indicates that these transcriptomic signatures reflect dynamic states associated with resistance development. We therefore asked whether these subtle baseline transcriptomic states are already present before treatment and associated with subsequent PARPi response in an independent pre-treatment cohort.

### Baseline Transcriptomic Signatures Recapitulate Longitudinal Patterns in an Independent PARPi-Treated Cohort

To investigate whether the subtle baseline transcriptomic differences between AR-pre and IR-pre tumors were reproducible, we applied the PRT and PST signatures defined in the longitudinal discovery cohort to an independent pre-treatment cohort of PARPi-treated ovarian cancer patients (n=71) (**Fig. 5a**). Initially, we stratified patients into AR (n=43) and IR (n=28) groups (**Methods**) and collected comprehensive clinical information to account for potential confounding factors in subsequent analysis (**Fig. 5a**). No statistically significant differences in clinical characteristics were observed between the AR and IR groups, although a modest difference in PARPi type distribution was noted (p = 0.079; **Supplementary Table 3**). We next examined whether these transcriptomic signatures differed according to BRCA mutation status. While BRCA-mutated (BRCAmt) tumors showed relatively uniform PRT and PST signature levels between the AR and IR groups (**Fig. 5b** and **c**), BRCA wild-type (BRCAwt) tumors exhibited significant differences in both PRT and PST signature levels (**Fig. 5d** and **e**). Specifically, BRCAwt IR tumors exhibited significantly higher PRT signature levels and significantly lower PST signature levels than AR tumors. Consistent with this stratified pattern, formal interaction analysis using the global PRT and PST signatures demonstrated significant effect modification by BRCA status for clinical response (interaction p = 0.04 and p = 0.007, respectively; **Supplementary Fig. 7a**). Pathway-specific analysis using cell-cycle PRT and HRR PST signatures largely recapitulated this pattern, with a significant AR–IR difference in cell-cycle PRT signature levels and a concordant trend in HRR PST signature levels within the BRCAwt subgroup (**Supplementary Fig. 7b**). These results demonstrate that the baseline transcriptomic differences distinguishing AR-pre and IR-pre tumors in our longitudinal discovery cohort were robustly recapitulated in an independent pre-treatment cohort, particularly within BRCAwt tumors. Notably, this recapitulation was most pronounced in BRCAwt tumors, where canonical BRCA-associated defects are absent.

**Figure 5.**
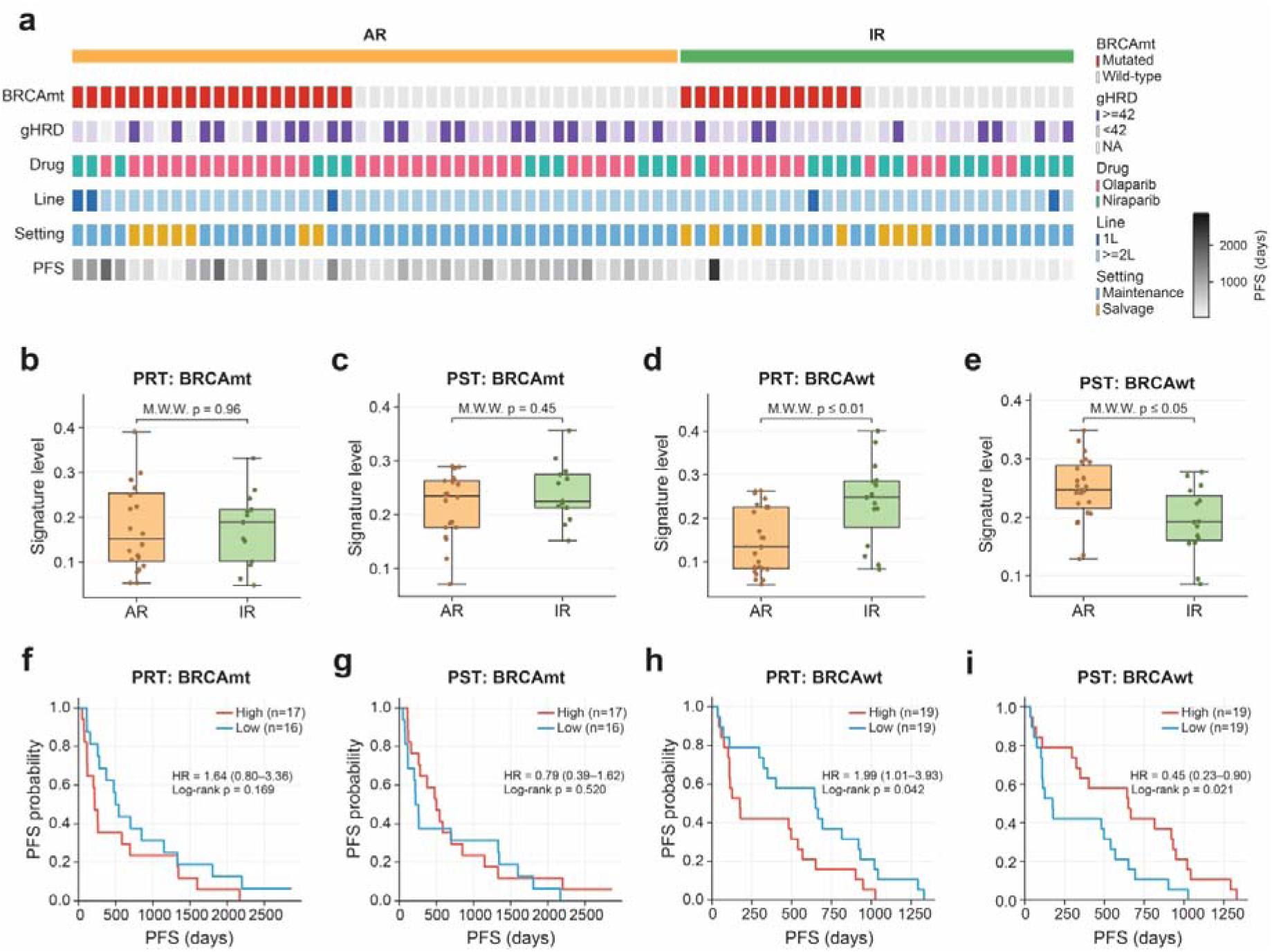
Validation of PRT and PST signatures in an independent pre-treatment PARPi cohort. (a) Clinical and molecular characteristics of the independent pre-treatment validation cohort. Each column represents an individual patient grouped by acquired resistance (AR) or innate resistance (IR). Annotations include BRCA mutation status, genomic homologous recombination deficiency (gHRD) status, administered PARPi types (olaparib, niraparib, or rucaparib), treatment line (1L or ≥2L), therapeutic setting (maintenance or salvage), and progression-free survival (PFS, days). (b, c) PRT and PST signature levels, respectively, in AR and IR tumors within the BRCA-mutated (BRCAmt) subgroup. P values were calculated using two-sided Mann-Whitney U tests. (d, e) PRT and PST signature levels, respectively, in AR and IR tumors within the BRCA wild-type (BRCAwt) subgroup. P values were calculated using two-sided Mann-Whitney U tests. (f, g) Kaplan–Meier analyses of PFS according to PRT and PST signature levels, respectively, within the BRCAmt subgroup. (h, i) Kaplan–Meier analyses of PFS according to PRT and PST signature levels, respectively, within the BRCAwt subgroup.

Next, to determine whether these transcriptomic signatures were also associated with clinical outcome, we evaluated their associations with PFS by stratifying the cohorts into high and low groups based on the median PRT or PST signature level. Within the BRCAmt group, neither transcript signature significantly stratified PFS (**Fig. 5f** and **g**). In contrast, within the BRCAwt group, high PRT signature levels were associated with significantly shorter PFS (log-rank p = 0.042), whereas low PST signature levels were associated with significantly shorter PFS (log-rank p = 0.021) (**Fig. 5h, I,** and **Methods**). A similar BRCA-stratified pattern was observed for the pathway-specific signatures, with cell-cycle PRT and HRR PST usage significantly associated with PFS in BRCAwt, but not BRCAmt, tumors (**Supplementary Fig. 7c**).

Collectively, these findings demonstrate that the transcriptomic signatures uncovered through longitudinal profiling are reproducible and remained detectable in pre-treatment tumor from an independent cohort, suggesting that adaptive transcriptomic programs may already exist at baseline. More importantly, the BRCA-dependent associations further suggest that PARPi resistance may arise through genetically constrained transcriptomic remodeling rather than a single universal mechanism. These results establish longitudinal isoform dynamics as a functional layer linking genetic background to adaptive drug resistance and provide a framework for understanding PARPi resistance beyond static genomic markers.

## Discussion

In this study, we demonstrate that longitudinal transcript-level analysis reveals mechanisms of PARP inhibitor (PARPi) resistance that are not captured by conventional gene-level expression profiling. While baseline clinical variables showed limited ability to explain progression-free survival (PFS) within the discovery cohort, dynamic changes in transcript usage (TU) consistently distinguished acquired resistance (AR) from innate resistance (IR). These findings suggest that transcript isoform regulation, rather than overall gene expression, represents a previously underappreciated layer of molecular adaptation during PARPi treatment.

A major finding of this study is that PARPi resistance is accompanied by coordinated TU remodeling in genes involved in homologous recombination (HR) and cell-cycle regulation. Importantly, these transcriptomic changes were not random but preferentially shifted toward functional isoforms capable of restoring DNA repair capacity, while reducing the usage of non-functional transcripts. This observation suggests a potential mechanism by which tumor cells can progressively regain DNA repair activity despite continued therapeutic pressure. Our results suggest that alternative splicing constitutes an additional adaptive mechanism capable of reshaping DNA repair pathways without requiring detectable changes at the DNA sequence or gene-expression level. Rather than acting as an independent process, transcript remodeling may function as a complementary regulatory layer that facilitates rapid functional adaptation under selective therapeutic pressure.

An additional observation is that IR tumors already exhibit a distinct transcriptomic landscape prior to treatment compared with AR tumors. Although the magnitude of these differences was modest, IR tumors consistently showed transcript usage patterns indicative of relatively higher DNA repair competency before treatment initiation. This suggests that PARPi responsiveness may not solely depend on the presence or absence of canonical genomic alterations but also on the pre-existing functional transcript landscape. From a mechanistic perspective, acquired resistance appears to represent a progressive remodeling process toward transcriptomic states that are already partially established in intrinsically resistant tumors, rather than a discrete molecular transition following treatment exposure.

Notably, the magnitude of transcriptomic remodeling during the development of acquired PARPi resistance was also associated with response to subsequent therapy. AR tumors from patients who subsequently experienced stable or progressive disease exhibited greater resistance-associated shifts in PRT and PST signature levels than those from patients who achieved partial response or complete response. This association suggests that the transcriptomic changes acquired during PARPi treatment may reflect a broader resistant state that extends beyond PARPi resistance itself. Given that the majority of subsequent therapies in this cohort were platinum-based (**Supplementary Table 2**), one possibility is that the transcriptomic adaptations accompanying acquired PARPi resistance may also be related to post-PARPi platinum cross-resistance. Although the mechanisms underlying response to subsequent therapies are likely heterogeneous, these findings raise the potential clinical value of resistance-associated transcriptomic remodeling as a biomarker not only of PARPi resistance but also of therapeutic susceptibility following PARPi failure.

Importantly, the transcriptomic signatures identified in the longitudinal cohort were independently validated in a separate pre-treatment cohort of patients who subsequently experienced recurrence. Patients stratified according to these PRT and PST signature levels revealed significant differences in progression-free survival, demonstrating that resistance-associated transcriptomic programs are detectable before treatment initiation and are related to the timing of recurrence. Furthermore, these associations were strongly influenced by BRCA mutation status. While transcript usage patterns effectively distinguish clinical response within specific genetic backgrounds, their contribution differed substantially between BRCA-mutant and BRCA-wildtype tumors. These findings suggest that the functional consequences of alternative splicing are highly dependent on the underlying genomic context. In tumors with existing defects in homologous recombination, transcript-level regulation may substantially influence the degree of residual DNA repair activity and subsequent therapeutic sensitivity. Conversely, in genetically distinct backgrounds, alternative regulatory mechanisms may dominate treatment response. Together, these observations highlight that genetic alterations and transcriptomic regulation should not be considered independent determinants of PARPi response but rather interacting layers that collectively define therapeutic susceptibility.

Despite these findings, our study also has several limitations. First, the transcript usage changes associated with PARPi resistance were not consistently driven by the same individual genes across patients. Instead, different tumors appeared to achieve similar functional outcomes through distinct combinations of transcript alterations. While this heterogeneity limits the identification of universal gene-level biomarkers, the reproducible enrichment of affected transcripts within common biological pathways, such as homologous recombination repair (HRR) and cell cycle, suggests that resistance develops through convergent functional adaptation rather than recurrent alterations in specific genes. Such diversity is consistent with the evolutionary nature of therapeutic resistance, where individual tumor clones may exploit multiple molecular routes to restore DNA repair capacity under selective pressure. Nevertheless, similar to the identification of recurrent hotspot mutations in cancer genomics, larger patient cohorts may eventually reveal recurrent transcript-level events that are currently underpowered to detect. Future studies with substantially expanded cohorts will therefore be essential to identify clinically actionable transcript biomarkers and determine whether recurrent isoform-switching events exist in PARPi-resistant disease.

Another important limitation arises from the use of short-read RNA sequencing. Transcript quantification and isoform assignment from short-read data inevitably introduce ambiguity, particularly for complex splicing events and highly similar transcript isoforms. To mitigate this limitation, our analyses focused on comparisons between functionally distinct transcript groups, such as protein-coding (Class 1) and predicted non-functional transcript (Class 3) classes, rather than relying on individual transcript estimates alone. We consider that this strategy substantially reduces the impact of transcript assignment errors while preserving biologically meaningful functional differences. Nevertheless, comprehensive characterization of resistance-associated splicing programs, precise identification of recurrent isoform switching events, and accurate reconstruction of full-length transcripts will ultimately require long-read RNA sequencing technologies capable of directly capturing complete transcript structures. Integration of longitudinal long-read transcriptomics with genomic profiling will therefore represent an important next step toward fully understanding the contribution of alternative splicing to PARPi resistance.

In summary, our study establishes transcript usage dynamics as an important molecular determinant of PARPi resistance that complements conventional genomic and gene-expression analyses. By revealing coordinated transcript remodeling associated with restoration of DNA repair capacity and demonstrating its prognostic relevance across independent patient cohorts, our findings highlight the value of transcript-level analyses in identifying biologically and clinically relevant changes in the context of both acquired and innate resistance, which are not readily captured by conventional gene-level expression analyses. More broadly, these results highlight alternative splicing as a clinically relevant layer of therapeutic adaptation and support future efforts to integrate transcript-resolved analyses into biomarker development and precision oncology.

## Methods

### Sequencing of cohort samples

RNA-seq data were generated from formalin-fixed/paraffin-embedded (FFPE) samples, RNA was extracted from 1 to 4 sections (4–10 μm thick) using a RNeasy FFPE mini-kit (Qiagen). RNA concentration was measured using a Qubit RNA HS Assay Kit (Thermo Fisher Scientific), and RNA quality was confirmed using a NanoDrop spectrophotometer (Thermo Fisher Scientific). RIN and DV200 values for RNA integrity were measured using a TapeStation 4200 system with RNA ScreenTape and Reagents (Agilent Technologies). For DV200 values < 40, ribosomal RNA (rRNA) was removed from RNA samples (500 ng) using an rRNA Depletion Kit (MGI Tech Co.). For DV200 values > 40, messenger RNA (mRNA) was isolated using a Dynabeads mRNA Purification Kit (Thermo Fisher Scientific). Library construction was performed using TruSeq RNA Library Prep kit v2 (Illumina inc., USA) or MGIEasy RNA Library Prep Set (MGI Tech Co.). Library concentration was measured using a Qubit dsDNA HS Assay Kit (Thermo Fisher Scientific), and library fragment size was measured using a TapeStation 4200 with D1000 ScreenTape and Reagents (Agilent Technologies). RNA paired-end sequencing with 100 bp reads was performed using Illumina Hiseq2500 sequencer and Hiseq SBS kit v4 (Illumina inc., USA), or DNBSEQ-G400 Sequencer and DNBSEQ-G400RS High-throughput Rapid Sequencing Kit (MGI Tech Co.). To align the raw reads of the discovery and validation samples, we trimmed using Trim Galore (v.0.6.5, https://www.bioinformatics.babraham.ac.uk/projects/trim_galore). The trimmed reads were aligned to the hg19 reference genome using a two-pass strategy with STAR (v.2.7.1a) with recommended options.

### Patient Cohort and Clinical Classification

Patients with advanced-stage or recurrent ovarian cancer who received PARP inhibitor (PARPi) as maintenance or salvage therapy at a single center between 2016 and 2021 were included. The longitudinal discovery cohort consisted of patients with paired tumor samples available before PARPi treatment (baseline) and after PARPi treatment (relapse). Pre-treatment samples were collected at the initial diagnosis before PARPi treatment, whereas post-treatment samples were obtained after progression on PARPi, either at secondary debulking surgery or biopsy. The validation cohort consisted of patients with only pre-PARPi samples. Relevant clinical information, including BRCA status, HRD status, PARPi type, line of therapy, treatment setting, progression status and time to progression on PARPi, subsequent therapy type, and progression on subsequent therapy, was obtained from electronic medical record review.

Patients were classified into Acquired Resistance (AR) and Innate Resistance (IR) groups based on treatment duration and objective response. For patients receiving PARPi as maintenance therapy, AR was defined as a treatment duration of ≥540 days (BRCA-mutated, first-line), ≥360 days (BRCA-mutated, second-line or later), ≥360 days (BRCA wild-type, first-line), or ≥180 days (BRCA wild-type, second-line or later). For patients receiving PARPi as salvage therapy, AR was defined as a best objective response of partial response (PR), complete response (CR), or no evidence of disease (NED). Patients not meeting these criteria were classified as IR. The discovery cohort comprised 40 patients (80 paired pre- and post-treatment samples). An independent validation cohort of 71 pre-treatment samples (43 AR and 28 IR) was used for external validation, classified using the same clinical criteria.

### Transcript Assembly and Quantification

RNA-seq reads in BAM format were processed using StringTie (v3.0.1)^33^. Individual transcript assemblies were generated for each of the 80 discovery cohort samples using the GENCODE v19 annotation as a reference guide (-p 4 -G gencode.v19.annotation.gtf)^34^. All per-sample GTF files were then merged into a unified consensus transcriptome using StringTie’s merge mode (--merge -c 5 -G gencode.v19.annotation.gtf), applying a minimum coverage threshold of 5 to exclude lowly supported transcripts. Among the assembled transcripts, 185,517 transcripts assigned to Ensembl-annotated protein-coding genes were retained for subsequent analyses.

Transcript-level quantification was subsequently performed for all samples in both the discovery and validation cohorts using StringTie’s estimation mode (-e -B -A gene_abund.tab), with the discovery-cohort merged GTF supplied as the reference annotation. Transcripts per million (TPM) values from the per-sample output GTF files were used as the final expression measure.

### Transcript Usage Calculation and DUT Identification

Transcript usage (TU) was defined as the proportional contribution of a given transcript to the total expression of its parent gene within a sample. For transcript t_i_ of gene g in sample s:

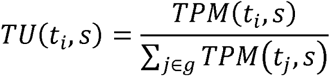

Differentially Used Transcripts (DUTs) were identified using paired Wilcoxon signed-rank tests on matched pre- and post-treatment TU values, applied separately within the AR and IR groups. Because multiple-testing correction markedly reduced statistical power given the large number of transcript-level tests and limited paired sample size, nominal p-values were used together with an effect-size threshold. Transcripts satisfying p < 0.05 and |ΔTU| ≥0.05 were defined as DUTs.

Before testing, lowly expressed transcripts were filtered by retaining those detected at TPM > 0 in at least 30% of samples, and only transcripts from genes without nominal evidence of post-treatment changes in total expression were included, as determined using the same paired Wilcoxon test (p ≥ 0.05), to focus on intra-gene transcript redistribution. A total of 161,621 transcripts remained after filtering and were used for DUT analysis.

#### Clinical classification

Among AR DUTs, transcripts with increased TU post-treatment (ΔTU > 0.05) were designated PARPi-resistance transcripts (PRTs), and those with decreased TU (ΔTU<−0.05) were designated PARPi-sensitivity transcripts (PSTs).

#### Functional classification

DUTs were further categorized into three classes. Class 1 (canonical protein-coding) comprised transcripts annotated as APPRIS Principal 1–5. Among the remaining transcripts^35^, those with a CPAT coding probability ≥0.364 (CPAT v3.0.5; recommended threshold for human transcripts)^36^ were assigned to Class 2 (non-canonical protein-coding), and those below this threshold were assigned to Class 3 (non-canonical non-coding).

### Differential Gene Expression Analysis

Differential gene expression analysis was performed separately within the AR and IR groups using DESeq2 in R^37^. Gene-level raw count matrices were used as input, and matched pre- and post-treatment samples were analyzed using a paired design that included patient identity and treatment condition (∼ patient + condition) to account for inter-patient variability. DEGs were defined as genes with an adjusted p-value <0.05 and |log2 fold change| >1. Volcano plots were generated using the EnhancedVolcano R package.

### Alternative Splicing Analysis

Alternative splicing (AS) events were quantified using SUPPA2 (v2.4)^38^. Events of seven types (skipped exon (SE), alternative 3′ splice site (A3), alternative 5′ splice site (A5), retained intron (RI), mutually exclusive exon (MX), alternative first exon (AF), and alternative last exon (AL)) were generated from the merged GTF using the generateEvents module. Per-sample percent spliced in (PSI) values were calculated with the psiPerEvent module in variable mode with a minimum read coverage of 10 (--mode variable --event-cov 10). Differential splicing between pre- and post-treatment was assessed separately within the AR and IR groups. For each of the seven event types, ΔPSI values were compared using the Mann–Whitney U test, and events with p < 0.05 and |ΔPSI| ≥0.1 were considered significant.

For splicing factor expression analysis, a curated list of 404 splicing factor (SF) genes was obtained from Seiler et al^39^. For each patient, treatment-associated expression changes were calculated across the curated SF genes and summarized as the mean ΔSF expression. Mean ΔSF expression values were then compared between the AR and IR groups using a two-sided Mann–Whitney U test.

### Cell Cycle Phase Scoring

Cell cycle phase activity was estimated computationally for each sample using gene set variation analysis (GSVA), implemented with the GSEApy package. Standard S-phase (43 genes) and G2/M-phase (54 genes) gene sets, derived from Seurat/Scanpy cell-cycle scoring marker lists, were used as input^40^. GSVA enrichment scores were calculated from gene-level TPM matrices using a Gaussian kernel (kcdf=’Gaussian’, min_size=10, max_size=500). Each sample was assigned a dominant cell cycle phase based on enrichment scores: samples with both S-phase and G2/M-phase scores ≤0 were classified as G1; among the remaining samples, those with S-phase score ≥ G2/M-phase score were classified as S-phase, and the rest as G2/M-phase. Phase distributions were then compared across AR and IR groups at pre- and post-treatment timepoints.

### Functional Enrichment Analysis

Pathway enrichment analysis was performed using the GSEApy package (Python)^41^ with the Reactome 2022 pathway database. Enrichment was conducted on genes associated with PRT-Class 1 and PST-Class 3 to characterize the biological pathways underlying each functional signature.

### Splicing-Based Scoring Systems

Two composite scores were developed to quantify the transcriptomic state of individual samples relative to resistance-associated isoform patterns. All TU values were z-score normalized prior to scoring.

#### Resistance-state score

Calculated as the difference between the mean z-scored TU of state-ordered PRTs and the mean z-scored TU of state-ordered PSTs:

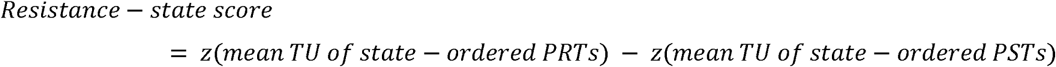

#### PARPi-pathway score

Calculated as the difference between the mean z-scored TU of cell-cycle PRTs and the mean z-scored TU of HRR PSTs:

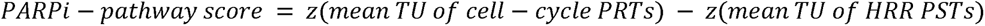

Both scores were evaluated across pre- and post-treatment timepoints in the discovery and in pre-treatment samples from the validation cohort, and their association with progression-free survival (PFS) was assessed.

### PCA and Projection onto the IR Axis

To quantify the longitudinal transcriptomic shift of AR patients toward an IR-like state, PCA and IR-axis projection were performed on the TU matrices of DUT features (PRT and/or PST sets).

#### Feature preparation

TU values were first transformed using the logit function with a small regularization constant ε = 10^−3^ to stabilize variance near the boundaries [0, 1]:

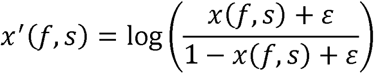

Transformed values were z-score normalized using the mean (μ_f̂{IR}) and standard deviation (σ_f̂{IR}) of IR pre-treatment samples:

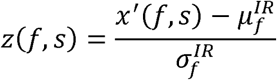

#### IR direction vector

The IR axis was defined as the unit vector pointing from the mean AR pre-treatment centroid to the mean IR pre-treatment centroid in z-score feature space:

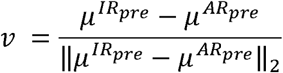

#### IR-axis projection score

Each sample *s* was projected onto the IR axis via the inner product:

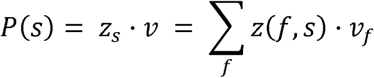

#### Longitudinal shift assessment

For each AR patient *p*, the treatment-induced change in projection score was computed as:

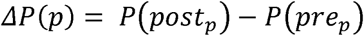

A one-sided Wilcoxon signed-rank test (alternative: ΔP > 0) was applied across AR patients to assess whether PARPi treatment significantly shifted tumors toward the IR transcriptomic state. The fraction of AR patients with ΔP > 0 was also reported.

#### PCA visualization

PCA (two principal components) was performed jointly on the concatenated pre- and post-treatment z-score matrices of all AR and IR samples. Trajectories were drawn connecting pre- and post-treatment coordinates of each AR patient, and a 1-σ covariance ellipse was overlaid on the combined IR sample coordinates.

### External Validation Cohort Analysis

To validate the generalizability of the identified signatures, the PRT and PST transcript lists derived from the discovery cohort were applied directly to an independent pre-treatment cohort (43 AR and 28 IR patients). TU values for these transcripts were computed from the validation cohort BAM files using the same StringTie quantification pipeline with the discovery-cohort merged GTF as reference. Splicing scores (Resistance-state score and PARPi-pathway score) were then calculated using the same definitions as in the discovery cohort, without any retraining or reselection of features. Between-group comparisons (AR vs. IR) were performed using the Mann-Whitney U test.

### Multivariable Cox Proportional Hazards Regression

The independent prognostic value of transcript-based signatures was evaluated using multivariable Cox proportional hazards (CoxPH) regression implemented with CoxPHFitter from the lifelines Python package. The primary outcome was progression-free interval (PFI), with event status coded as a binary variable (0 = censored, 1 = event). Samples with missing values in PFI, survival status, or any covariate were excluded. Covariates with fewer than two unique non-missing values were also excluded. Results were reported as hazard ratios (HR) with 95% confidence intervals and p-values, and visualized as forest plots.

### Statistical Analysis and Visualization

All statistical analyses and visualizations were implemented in Python. In the discovery cohort, paired pre-vs. post-treatment comparisons were performed using the Wilcoxon signed-rank test, with p-values annotated using the stat annotations package. Between-group comparisons in the validation cohort were performed using the Mann-Whitney U test. Kaplan-Meier survival curves were compared using the log-rank test. All plots were generated with the seaborn library.

## Supplementary Information

## Supplementary data

Supplementary Data are available at Supplemental Material online. Raw RNA sequencing data generated from our cohort samples have been deposited in the SRA (reviewer access: PRJNA1532479).

## Ethics approval and consent to participate

Not applicable.

## Author contributions

**J. Yoo:** Conceptualization, data curation, formal analysis, investigation, visualization, methodology, writing – original draft. **Y.-N. Kim:** Conceptualization, data curation, formal analysis, investigation, writing – review & editing. **Y. Jung:** Data curation, formal analysis. **E.-H. Cho:** Resources, data curation. **J.K. Choi:** Supervision, funding acquisition, project administration. **H.G. Kang:** Conceptualization, data curation, formal analysis, investigation, supervision, writing – original draft. **J.-Y. Lee:** Conceptualization, resources, data curation, formal analysis, supervision, project administration.

