## Supplementary Figures for "Longitudinal transcriptomic analysis reveals adaptive transcript usage remodeling associated with PARP inhibitor resistance"

**Supplementary Figure 1.** Pathway enrichment analysis of differentially expressed genes in AR and IR tumors

**Supplementary Figure 2.** Transcript usage changes in PARPi-related genes in AR tumors

**Supplementary Figure 3.** Patient-level heterogeneity of Class1 PRTs and Class3 PSTs Usage changes in AR tumors

**Supplementary Figure 4.** Pathway-specific PRT and PST patterns across PARPi resistance states

**Supplementary Figure 5.** Association of pathway-specific transcriptomic remodeling with subsequent treatment outcomes

**Supplementary Figure 6.** Gene-level examples of pathway-specific transcript usage remodeling across PARPi resistance states

**Supplementary Figure 7.** BRCA-dependent associations of PRT and PST signatures in the independent validation cohort

**Supplementary Tables**

**Supplementary Table 1.** Clinical characteristics of the longitudinal discovery cohort

**Supplementary Table 2.** Subsequent treatment regimens in patients with acquired PARPi resistance

**Supplementary Table 3.** Clinical characteristics of the independent pre-treatment validation cohort

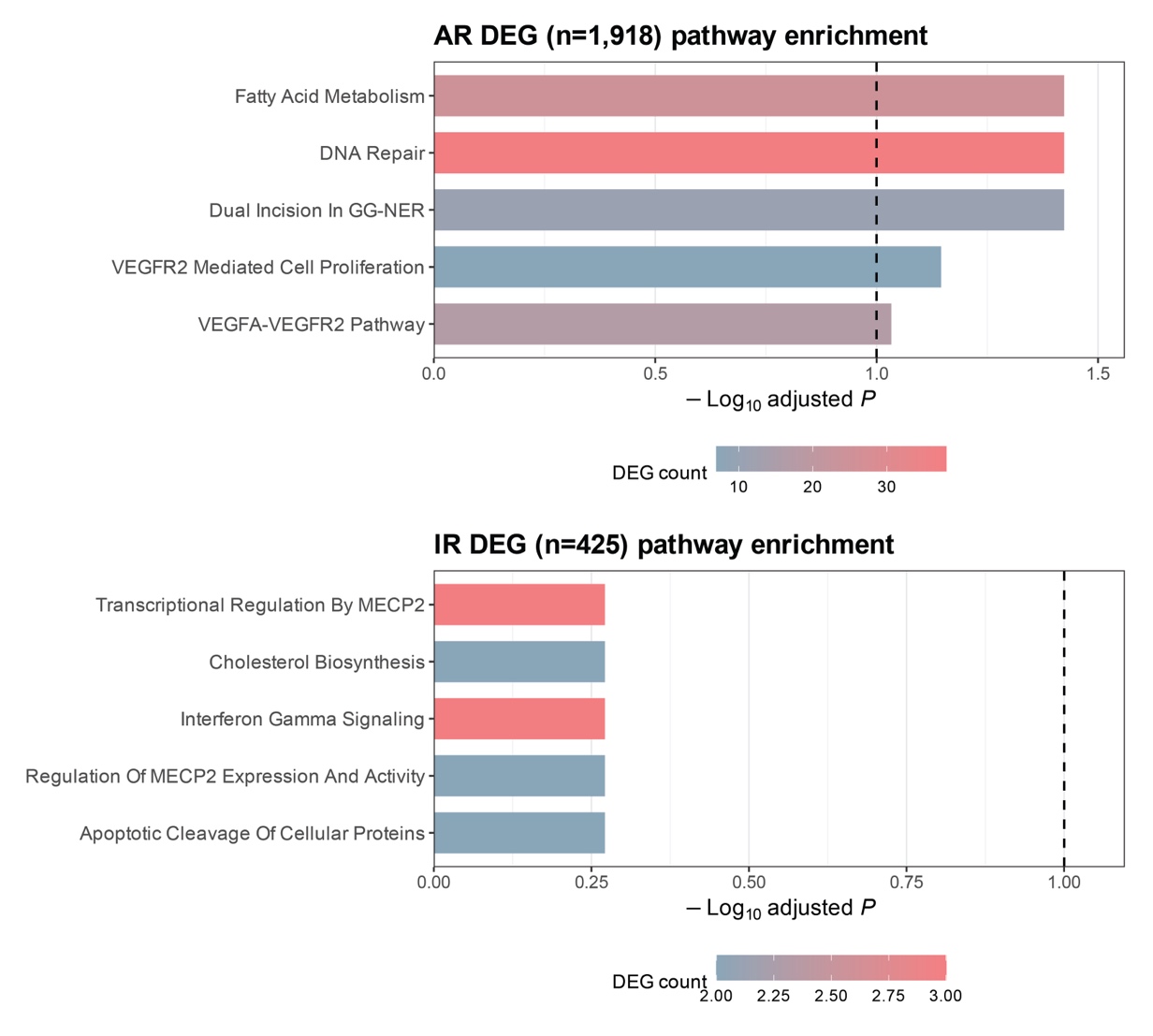

**Supplementary Figure 1. Pathway enrichment analysis of differentially expressed genes in AR and IR tumors.**

Pathway enrichment analysis of differentially expressed genes (DEGs) identified from paired pre- and post-treatment comparisons in the AR (n=1,918 DEGs) and IR (n=425 DEGs) groups. Bars represent -log10 adjusted P values, and colors indicate the number of DEGs associated with each pathway. The dashed line indicates FDR = 0.1. Significant enrichment of DNA repair-related pathways was observed in AR but not IR tumors.

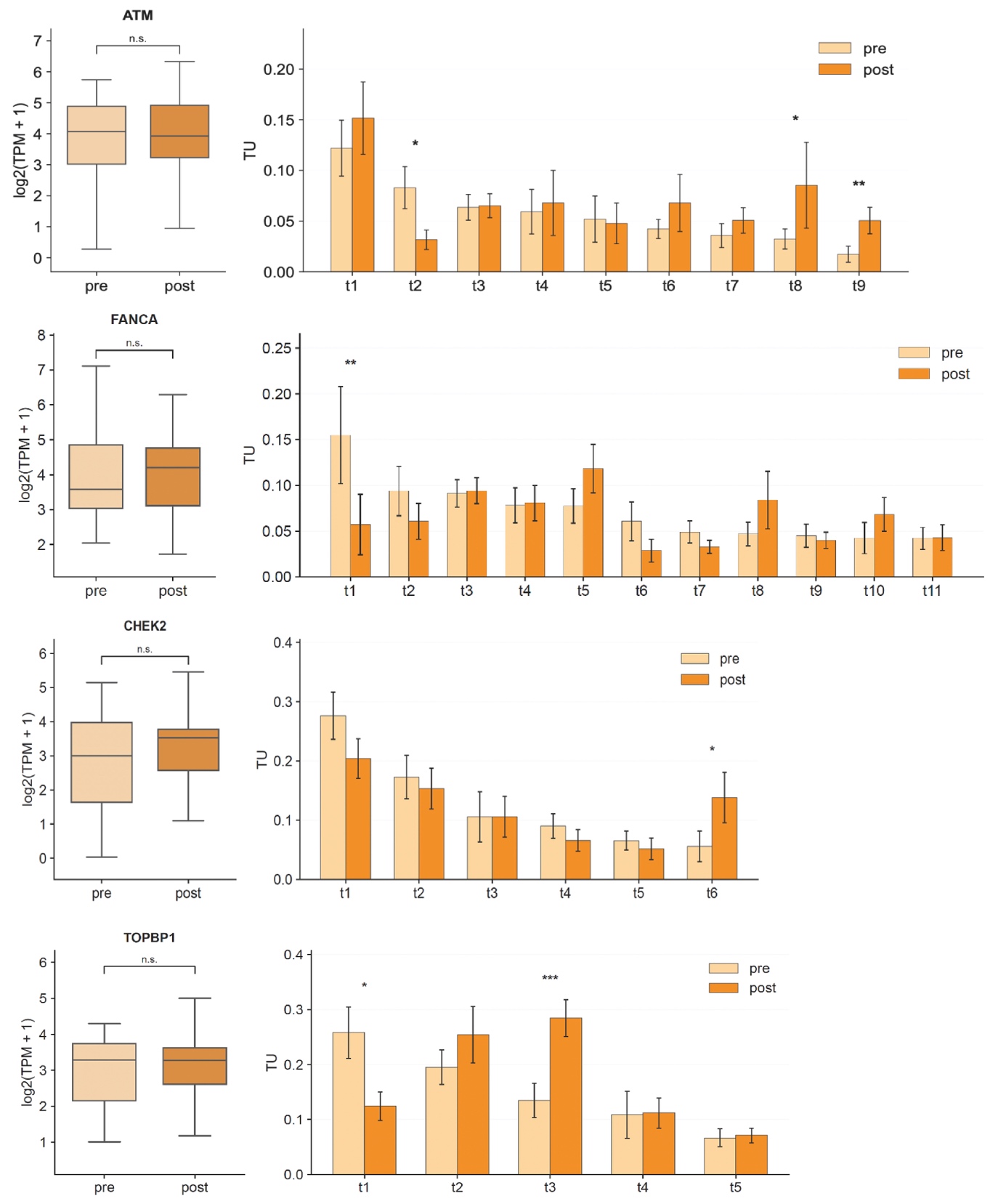

**Supplementary Figure 2. Transcript usage changes in PARPi-related genes in AR tumors.**

Representative PARPi-related genes showing transcript usage changes between paired pre- and post-treatment AR tumors despite limited changes in total gene expression. Gene-level expression (left) and transcript-level TU (right) are shown for ATM, FANCA, CHEK2, and TOPBP1. ‘t1’-‘tn’ denote individual transcript isoforms. P values were calculated using two-sided Wilcoxon signed-rank tests.

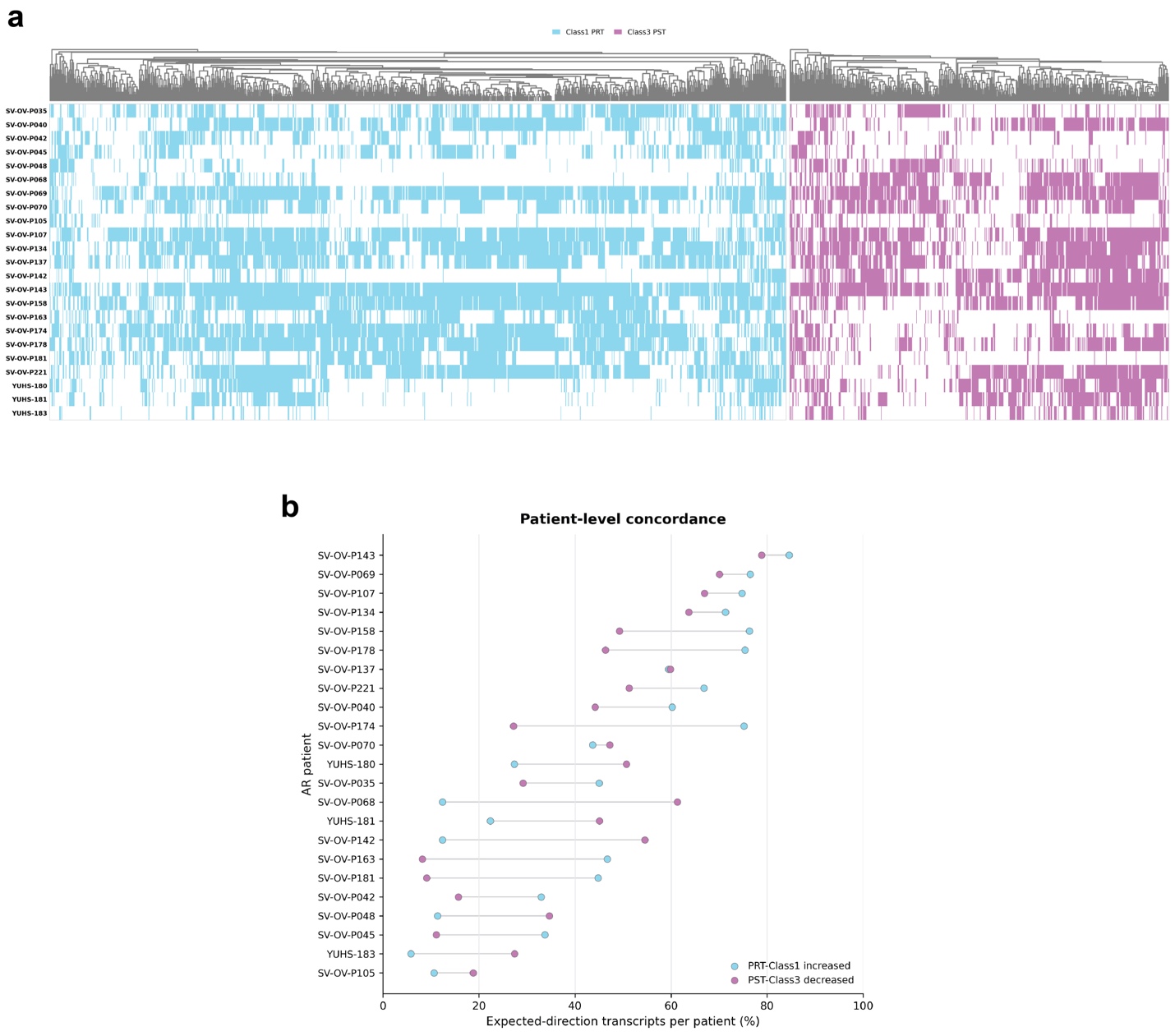

**Supplementary Figure 3. Patient-level heterogeneity of Class1 PRTs and Class3 PSTs Usage changes in AR tumors.**

(a) Patient-by-transcript heatmap showing treatment-associated changes in Class1 PRTs and Class3 PSTs across individual AR tumors. Colored cells indicate significant transcript usage changes in the expected resistance-associated direction (increased TU for Class1 PRTs and decreased TU for Class3 PSTs), whereas uncolored cells indicate transcripts not meeting these criteria.

(b) Patient-level proportions of Class1 PRTs showing increased TU and Claass3 PSTs showing decreased TU following treatment.

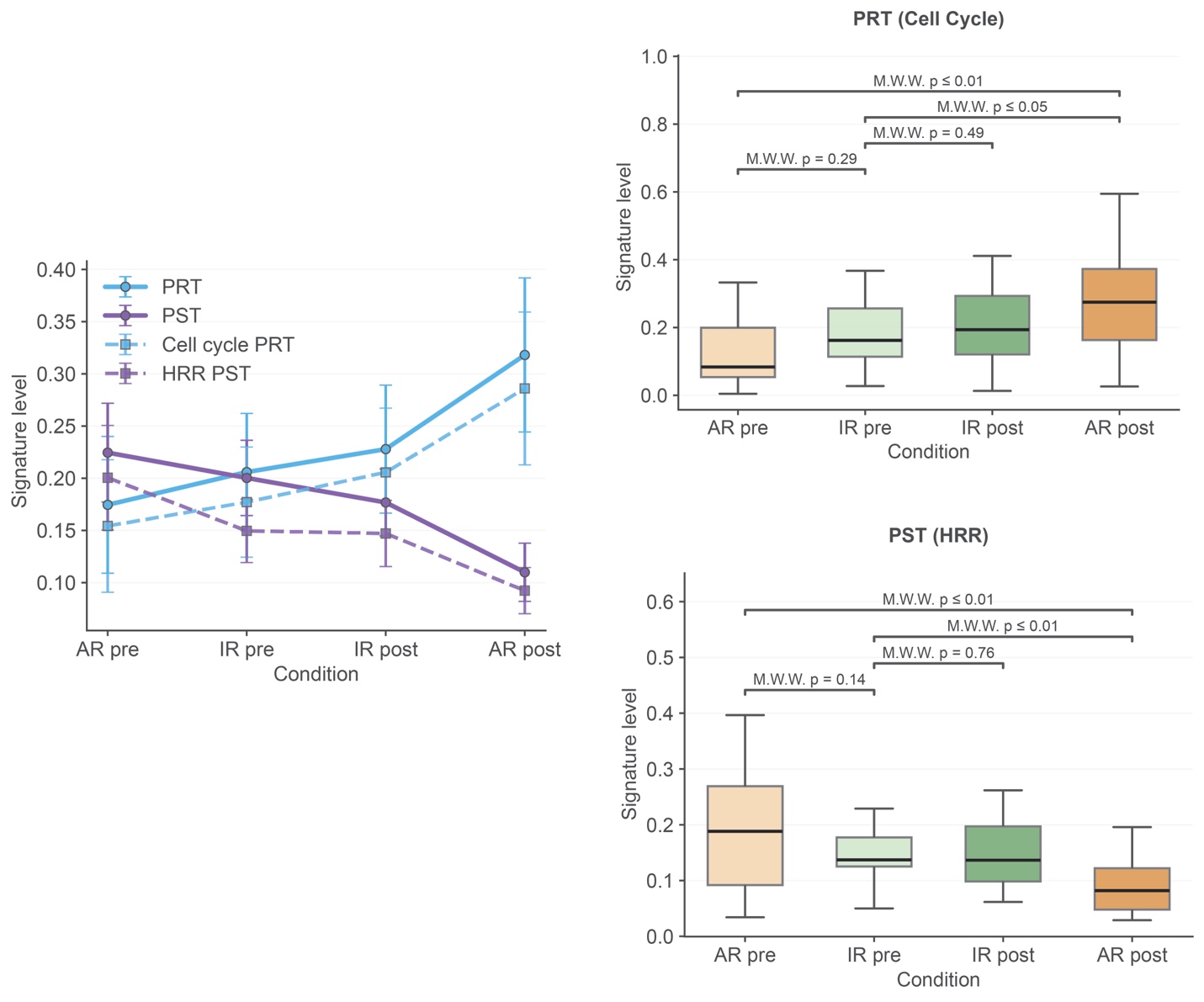

**Supplementary Figure 4. Pathway-specific PRT and PST patterns across PARPi resistance states.**

Comparison of global PRT/PST signatures and pathway-specific cell-cycle PRT and HRR PST signatures across AR pre-, IR pre-, IR post-, and AR post-treatment tumors. Pathway-specific signatures showed resistance-associated patterns consistent with the global PRT/PST signatures. P values were calculated using two-sided Mann-Whitney U tests for comparisons between independent groups and Wilcoxon signed-rank tests for paired pre- and post-treatment comparisons.

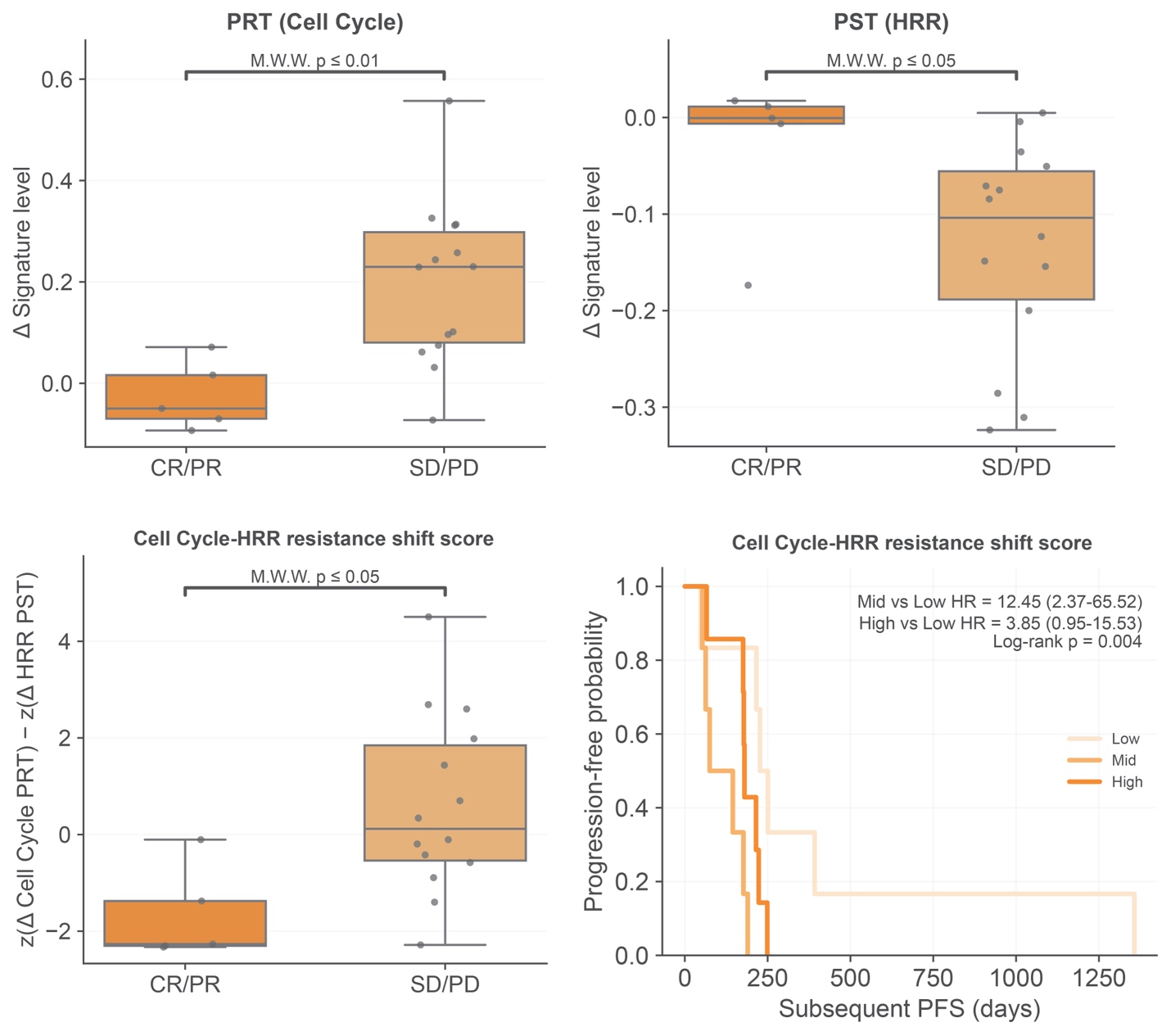

**Supplementary Figure 5. Association of pathway-specific transcriptomic remodeling with subsequent treatment outcomes.**

Changes in cell-cycle PRT and HRR PST signature levels during PARPi treatment according to subsequent treatment response among AR tumors. The Cell Cycle-HRR resistance shift score integrates opposing changes in cell-cycle PRT and HRR PST signature levels. Kaplan-Meier curves show subsequent PFS according to tertiles of the pathway-specific resistance shift score. P values for CR/PR versus SD/PD comparisons were calculated using two-sided Mann-Whitney U tests; survival distributions were compared using the log-rank test, with hazard ratios estimated using Cox proportional hazards models.

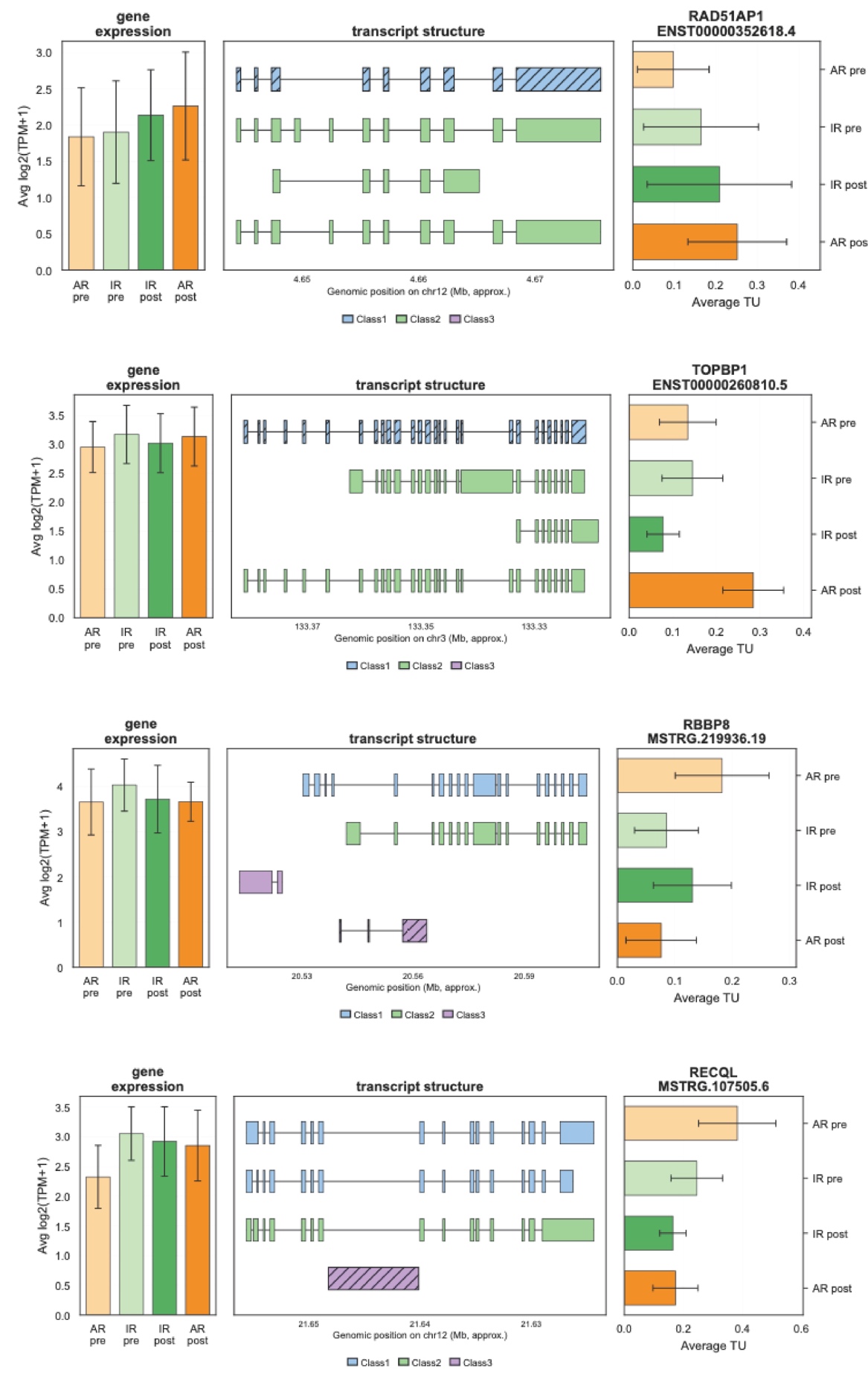

**Supplementary Figure 6. Gene-level examples of pathway-specific transcript usage remodeling across PARPi resistance states.**

Gene expression, transcript structures, and transcript usage of representative cell-cycle PRTs (*RAD51AP1* and *TOPBP1*) and HRR PSTs (*RBBP8* and *RECQL*) across the four clinical groups. Transcript structures are annotated according to functional class, and bar plots show the mean TU of each representative transcript.

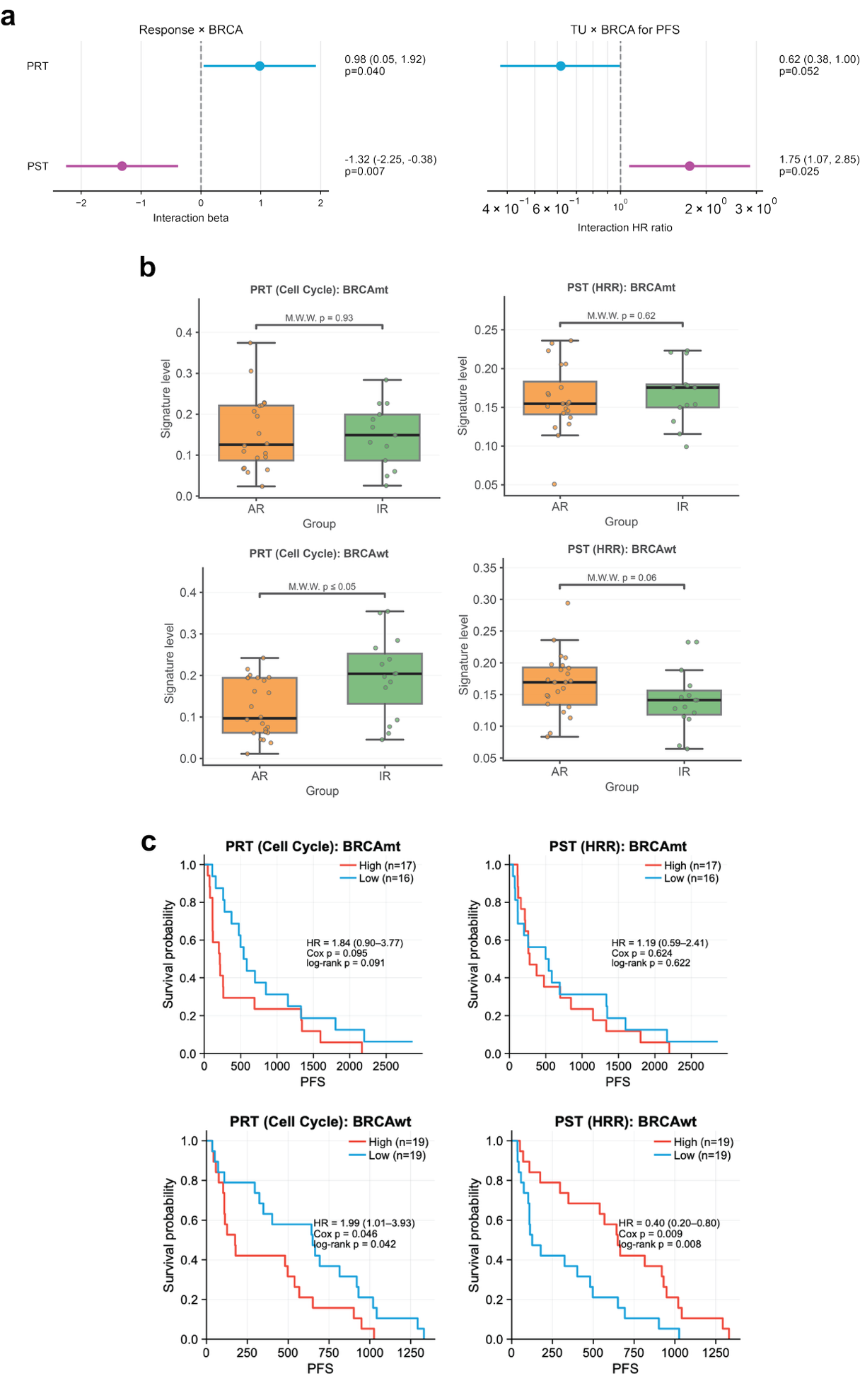

**Supplementary Figure 7. BRCA-dependent associations of PRT and PST signatures in the independent validation cohort.**

(a) Interaction analyses evaluating modification by BRCA mutation status of the associations between global PRT/PST signature levels and clinical response (left) or PFS (right). Points and error bars indicate interaction β coefficients (left) or ratios of hazard ratios (right) and their 95% confidence intervals. P values are from tests of the clinical response × BRCA mutation status interaction term in ordinary least squares (OLS) regression models (left) and the signature level × BRCA mutation status interaction term in Cox proportional hazards regression models (right).

(b) Mean cell-cycle PRT and HRR PST signature levels in AR and IR tumors stratified by BRCA mutation status. P values were calculated using two-sided Mann-Whitney U tests.

(c) Kaplan-Meier analyses of PFS according to cell-cycle PRT and HRR PST signatures, stratified by BRCA mutation status. Patients were classified into high- and low-usage groups using median TU. Hazard ratios were estimated using Cox proportional hazards models, and survival distributions were compared using log-rank tests.
